# Epigenetic inactivation of miR-203 as a key step in neural crest epithelial-to-mesenchymal transition

**DOI:** 10.1101/392142

**Authors:** Estefanía Sánchez-Vásquez, Marianne E. Bronner, Pablo H. Strobl-Mazzulla

**Author notes:** Correspondence should be addressed to PHS-M.

## Abstract

miR-203 is a tumor-suppressor microRNA with known functions in cancer metastasis. Here, we explore its normal developmental role in the context of neural crest development. As neural crest cells undergo an epithelial-to-mesenchymal transition to emigrate from the neural tube, miR-203 displays a reciprocal expression pattern with key regulators of neural crest delamination, *Phf12* and *Snail2,* and interacts with their 3’UTRs. Ectopic maintenance of miR-203 inhibits neural crest migration, whereas its functional inhibition using a “sponge” vector promotes premature neural crest delamination. Bisulfite sequencing further shows that epigenetic repression of miR-203 is mediated by the *de novo* DNA methyltransferase DNMT3B, whose recruitment to regulatory regions on the miR-203 locus is directed by SNAIL2 in a negative feedback loop. These findings reveal an important role for miR-203 in an epigenetic-microRNA regulatory network that influences the timing of neural crest delamination.

**Summary statement:** The EMT is a highly conserved process, involving similar levels of regulation in both neural crest and cancer cells. Our work shows an epigenetic-miRNA-gene regulatory circuit, conserved in cancer, which controls the timing of neural crest EMT as well.

## Introduction

Neural crest cells (NCC) are a transient embryonic cell population that arise in the neuroectoderm, then migrate to the periphery where they contribute to diverse derivatives including craniofacial bone and cartilage, neurons and glia of the peripheral nervous system, pigment cells, and portions of the cardiovascular system (Crane and Trainor, 2006). NCC emigrate from the forming central nervous system by undergoing an epithelial-mesenchymal transition (EMT), similar to that observed during initiation of tumor metastasis (Kerosuo and Bronner-Fraser, 2012). An evolutionarily conserved gene regulatory network (GRN) regulates NCC EMT and depends upon the coordinated action of transcription factors including *Snail1/2, Zeb2 (Sip1),* and *FoxD3* (Simoes-Costa and Bronner, 2015). In addition to transcriptional regulators mediating neural crest EMT, epigenetic mechanisms including DNA methylation (Hu et al., 2012, Hu et al., 2014) and histone modifications (Strobl-Mazzulla and Bronner, 2012, Strobl-Mazzulla et al., 2010) are also at play. In particular, a repressor complex, comprised of SNAIL2 and the epigenetic reader PHF12 represses *cadherin 6b (Cad6b)* (Strobl-Mazzulla and Bronner, 2012), whose down-regulation is required for initiation of neural crest delamination from the neural tube. These results demonstrate an additional level of fine tuning and important role for epigenetic regulators during neural crest EMT, raising the intriguing possibility that other yet to be identified factors may be involved.

In recent years, microRNAs (miRNAs) have been shown to be key regulators of EMT in several tumor cells (Diaz-Lopez et al., 2014, Ding, 2014, Xia and Hui, 2012). MicroRNAs are ~22-nucleotides single-stranded RNAs that negatively regulate gene expression post-transcriptionally (Kloosterman and Plasterk, 2006) by inhibiting translation and/or causing degradation by binding to complementary sequences located at the 3’-untranslated region (3’-UTR) of target mRNAs. Approximately 50% of miRNAs genes are embedded or associated with CpGs islands, and their expression is most often regulated by methylation of cytosines therein (Weber et al., 2007). In several tumor cells, it has been shown that hypermethylation of anti-tumoral miRNAs leads to initiation of EMT (Ahmad et al., 2014, Kiesslich et al., 2013, Lujambio et al., 2008, Wang et al., 2012, Bonnomet et al., 2010, Diaz-Lopez et al., 2014, Ding, 2014, Nelson and Weiss, 2008, Xia and Hui, 2012). The commonalities between the migration of cancer cells and embryonic neural crest cells (Mayor and Theveneau, 2013, Scarpa and Mayor, 2016, Friedl and Gilmour, 2009) suggest the intriguing possibility that similar epigenetic-microRNA to those functioning in metastasis may be involved in NCC development. As case in point, we describe in avian embryos that the epigenetic repression of miR-203, directed by SNAIL2 enables upregulation of two of its direct targets, *Phf12* and *Snail2,* necessary for the delamination of the neural crest from the neural tube. These findings support the idea that a single microRNA, regulating key genes of the EMT process, may “throw” a precise switch mediating an epithelial to a mesenchymal state of neural crest cells.

## Results

### miR-203 expression is reduced at the time of NCC delamination

Given that *Phf12* and *Snail2* are involved in regulation of NCC EMT (Strobl-Mazzulla and Bronner, 2012), we performed an *in silico* and literature analysis to investigate miRNAs that might regulate these transcription factors. Interestingly, we found 9 and 7 families, respectively, of miRNAs in the 3’UTRs of *Phf12* and *Snail2* with sites conserved across vertebrates (Table S1). Based on its their reported functionality, demonstrated target genes and expression pattern during chick development (Darnell et al., 2006), we further focused on miR-203 for in depth analysis. Importantly, miR-203 has been described to act as tumor suppressor (Benaich et al., 2014, Miao et al., 2014, Zhu et al., 2013b), that directly regulates *Snail2* expression (Gao et al., 2017, Shi et al., 2015, Zhang et al., 2015, Xiao et al., 2017), whose epigenetic repression causes metastasis in several tumor cells including NCC-derived melanoma (Boldrup et al., 2012, Boll et al., 2013, Chen et al., 2012, Chiang et al., 2011, Ding et al., 2013, Furuta et al., 2010, Huang et al., 2014, Ju et al., 2014, Moes et al., 2012, Zhang et al., 2014, Zhao et al., 2013, Bu and Yang, 2014, Lohcharoenkal et al., 2018). Moreover, the mature miR-203 sequence is highly conserved throughout vertebrates including the basal lamprey (Fig. 1B), suggesting an ancient and conserved function.

**Figure 1:**
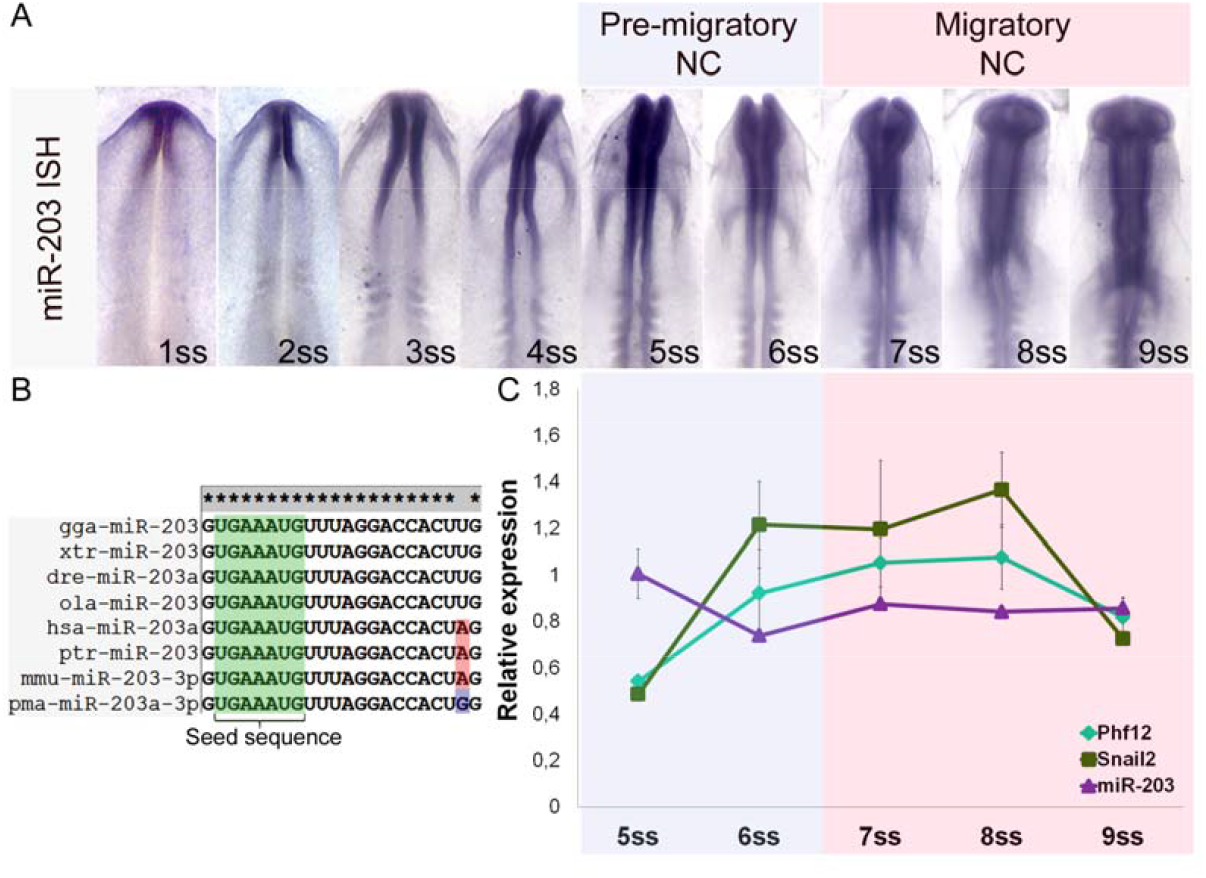
miR-203 expression is reduced at the time of NCC delamination. (A) Whole-mount *in situ* hybridization (ISH) using LNA-DIG labelled probes reveals a specific expression of the mature miR-203 on the neural tube which is reduced from 5 somite stage (ss) to 9ss. (B) Conservation analysis of miR-203 in vertebrates. Gga *(Gallus gallus),* xtr (Xenopus tropicalis), dre *(Danio* rerio), ola *(Oryzias* latipes), hsa *(Homo* sapiens), ptr *(Pan* troflodytes), mmu *(Mus* musculus), pma *(Petromyzon marinus).* (C) RT-qPCR analyses show reducing miR-203 expression from 5 to 6ss in an opposite manner than the observed increase on *Snail2* and *Phf12* expression at the beginning of NCC delamination.

As a first step in analyzing its possible function during NCC development, we examined the expression pattern of miR-203 transcripts by *in situ* hybridization (ISH) in early chick embryos. By using LNA-DIG labelled probes, we found that mature miR-203 expression begins during gastrulation by stage 4 (Fig. S1A). During neurulation at the 1-4 somite stage (ss), miR-203 is consistently expressed in the forming neural tube (Fig. 1A). Interestingly, we observed a clear reduction by the 5 to 8ss in the miR-203 expression in the cranial neural tube, corresponding to the initiation of neural crest emigration. Analyses by stem-loop RT-qPCR confirmed that mature miR-203 expression is reduced from 5 to 6ss, coincident with the increase of *Snail2* and *Phf12* expression at the time of NCC delamination (Fig. 1C). These results are consistent with the intriguing possibility that miR-203 has an important role in neural crest EMT.

### miR-203 locus is highly methylated on pre-migratory NCC

In several metastatic tumor cells, the miR-203 locus is epigenetically silenced by DNA methylation (Chim et al., 2011a, Chim et al., 2011b, Diao et al., 2014, Furuta et al., 2010, Huang et al., 2014, Taube et al., 2013, Zhang et al., 2013, Zhang et al., 2011). Interestingly, we found by *in silico* analysis that the miR-203 locus is embedded in a CpG island (Fig. 2A). On this basis, we selected two genomic regions, one at the putative proximal promoter (region 1) and the other at the beginning of the pri-miR-203 (region 2), to analyze the DNA methylation abundance on pre-migratory (PM-NCC) or migratory (M-NCC) neural crest cells and the ventral neural tube (NT). By using bisulfite conversion, we observed high enrichment on methylated CpGs in region 2, but not on region 1, in the PM-NCC compared with the low abundance detected on the M-NCC and NT (Fig. 2B-C). These significant differences are clearly evident when comparing the total percentage of methylated CpGs (Fig. 2D-E) from the different samples. We identified that 18.4% and 28.9% of the CpGs are methylated on region 1 and 2 on PM-NCC, compared with 9.5% and 6.1% on the NT, and 5.3% and 7% detected on the M-NCC, respectively. These results suggest that the decreased expression of miR-203 in pre-migratory NC may be the consequence of hypermethylation of its locus.

**Figure 2:**
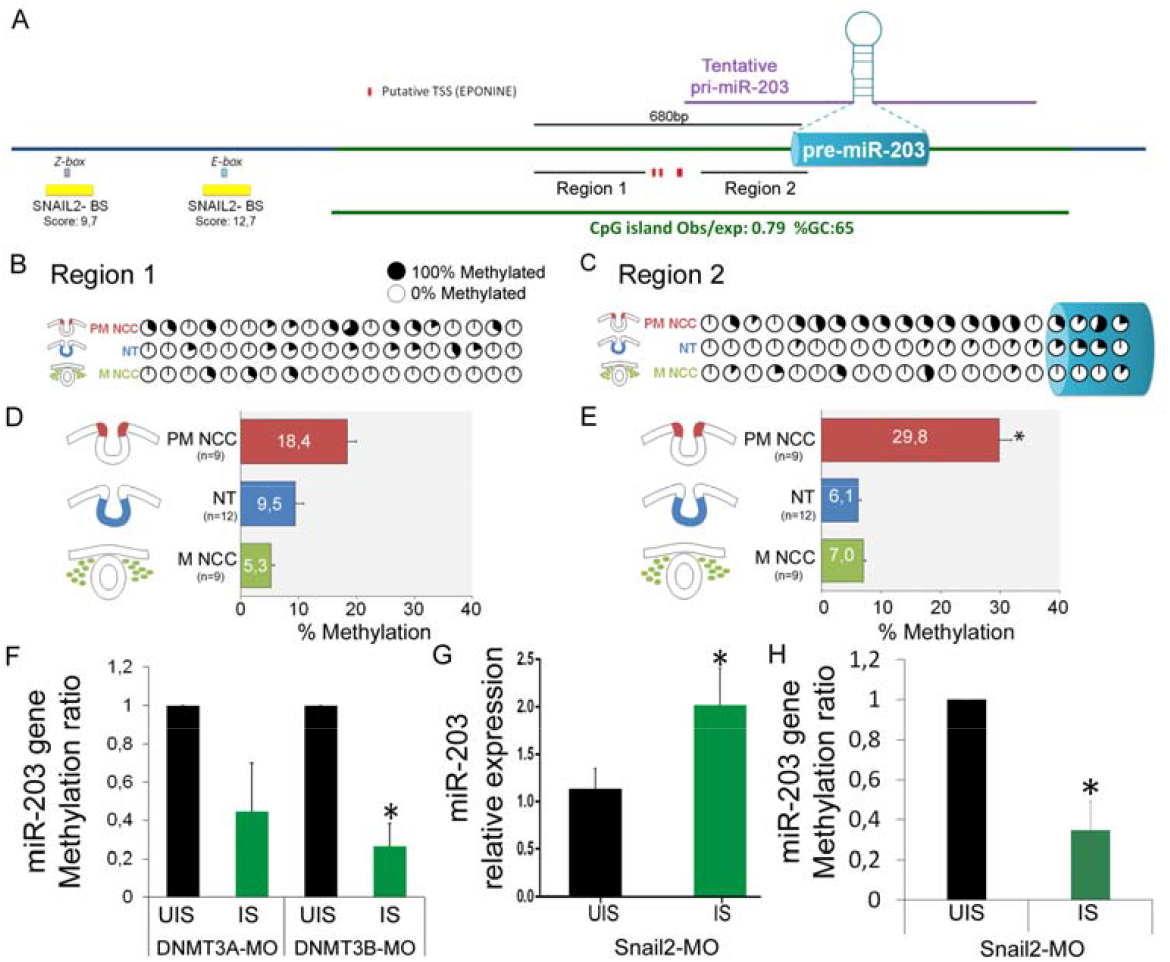
SNAIL2 and DNMT3B are required for DNA methylation on miR-203 locus on premigratory NCC. (A) miR-203 genomic context. The CpG island content (green bar) for miR-203 gene was identified using the UCSC genome browser (https://genome.ucsc.edu/) and Ensembl (http://www.ensembl.org/). The putative transcriptional start sites (TSS) were obtained using Eponine (https://www.sanger.ac.uk/science/tools/eponine). The putative binding sites for SNAIL2 were mapped using JASPAR 2018 (http://jaspar.genereg.net/). We also show the two regions used in our bisulfite sequencing and the pre-miR-203 accordingly to the miRbase (http://www.mirbase.org/). Bisulfite sequencing profiles of CpGs methylation on the region 1 (B) and region 2 (C) were analyzed on pre-migratory NCC (PM-NCC), migratory NCC (M-NC), and ventral neural tube (NT). Percentage of each methylated CpG sites are shown with filled (100% methylated) and open (0% methylated) circles. Bar graph represent the total percentages of methylated CpGs on the different regions for the three analyzed samples on the region 1 (D) and the region 2 (E). We evidenced in PM-NCC a higher percentage of methylated CpGs compared with the other samples. Number in brackets indicated the analyzed sequences. Asterisk (*) indicate significant differences by ANOVA. (F) Morpholino-mediated loss of DNMT3A (DNMT3A-MO) and DNMT3B (DNMT3B-MO) function results in a reduction of methylated CpGs on the injected side (IS) compared with the uninjected side (UIS) of the same group of embryos. Morpholino-mediated loss of SNAIL2 (SNAIL2-MO) function maintains an elevated level of miR-203 expression (G) and reduced CpGs methylation on the region 2 (H) on the IS compared with the UIS of the same group of embryos. Asterisk (*) indicate significant differences by Student’s t-test.

### DNMT3B and SNAIL2 are required for miR-203 methylation

De novo DNA methyltransferases DNMT3A and 3B are both involved in neural crest development (Hu et al., 2012). To examine the possibility that DNA methylation may be involved in regulating miR-203 expression, we examined the effects of loss of DNMT3A and 3B on miR-203 expression. To this end, we performed bisulfite sequencing after loss of function experiments with previously characterized fluorescein-tagged morpholino oligonucleotides, DNMT3A-MO and DNMT3B-MO (Hu et al., 2012, Hu et al., 2014). After unilateral injection and electroporation, dorsal neural tubes from 6ss embryos were dissected and bisulfite converted to analyze the methylation abundance on region 2 of miR-203 locus (Fig. 2A). The results show that loss of function of either of the two DNMTs consistently decreases the abundance of methylated CpGs in the region 2 (Fig. 2F). However, depletion of DNMT3B, but not DNMT3A, resulted in significant differences in methylation of miR-203.

Transcription factors specifically bind to target DNAs to recruit and guide DNA methyltransferases to specific genomic sites (Siegfried and Simon, 2010). During tumor metastasis, SNAIL2 binds to the miR-203 promoter to inhibit its transcription (Ding et al., 2013), though the mechanism underlying this repression is unknown. Our bioinformatic analysis reveal several SNAIL2-binding sites ~1kb upstream of the pre-miR-203 (Fig. 2A, Table S2), two of which have a high binding score (>9) accordingly to JASPAR 2018 (http://jaspar.genereg.net/). To test the effects of SNAIL2 loss of function, we electroporated a previously characterized Snail2 morpholino (Snail2-MO) (Taneyhill et al., 2007) unilaterally and dissected half dorsal neural tubes from 6ss embryos for analysis of miR-203 expression and DNA methylation. Interestingly, the results show that Snail2 knockdown causes a significant upregulation of miR-203 expression (Fig. 2G). Consistent with this finding, we found a significant reduction in the abundance of methylated CpGs in region 2 of the miR-203 locus (Fig. 2H). These results suggest that SNAIL2 is involved in the epigenetic repression of miR-203 in the premigratory neural crest.

### Overexpression of miR-203 causes loss of migrating NCC

Since miR-203 expression is epigenetically repressed at the initiation of their migration, we asked whether maintenance of miR-203 expression would prevent neural crest delamination. To this end, we designed an overexpression vector in which we cloned the pre-miR-203 sequence, plus a few hundred base pair arms for correct processing, under the control of CAG promoter (pCAG-203) (Fig. 3A). Overexpression of miR-203 in pCAG-miR-203 electroporated embryos was confirmed by LNA-ISH (Fig. S2B). To demonstrate that the pCAG-miR-203 expresses a functional miR-203, we first used a two-colored sensor vector (Cao et al., 2007) comprised of a nuclear-localized destabilized EGFP with a half-life of 4 hours (d4EGFP_N_), driven by a CAG promoter, followed by a 3’UTR containing two copies of a bulged complementary sequence for miR-203. In addition, the vector contains a nuclear-localized monomeric red fluorescent protein (mRFPN) driven by another CAG promoter that serves as an electroporation control (see scheme of pSdmiR-203 in figure S2C). Together with the dual sensor pSdmiR-203, we co-electroporated either control empty vector (left) or the vector expressing miR-203 (right) onto each side of the embryo (Fig. S2D). The results show that on the control side, most of the electroporated cells appeared yellow, due to expression of d4EGFPN and mRFP_N_. However, on the experimental side, overexpression of miR-203 causes a strong reduction of d4EGFPN and the cells only express mRFPN (Fig. S2E). These results confirm the functionality of our miR-203 expression construct. Gain-of-function experiments were conducted by electroporating the miR-203 vector on one side of the embryo (Fig. S2A). The results show that miR-203 overexpression causes a drastic reduction in the numbers of migratory neural crest cells identified by *Sox10* expression (Fig. 3B). By categorizing embryos according to their phenotype (normal or reduced migratory NCC), we observed a significant increase in the number of embryos with a reduced number of migratory NCC on pCAG-203 electroporated side compared with control embryos electroporated with an empty vector (pCAG) (Fig. 3C). To rule out the possibility that this may be secondary to a specification defect, we analyzed the expression of early neural crest specifier genes by immunohistochemistry at pre-migratory stages. Interestingly, our results demonstrate that overexpression of miR-203 does not affect expression of FOXD3, one of earliest neural crest specification markers (Fig. 3D), suggesting that specification occurs normally. In contrast, there was a clear reduction in SNAIL2, demonstrating reciprocal repression. Taken together, our results highlight the role of miR-203 in NCC delamination, likely by affecting EMT inducers like SNAIL2.

**Figure 3:**
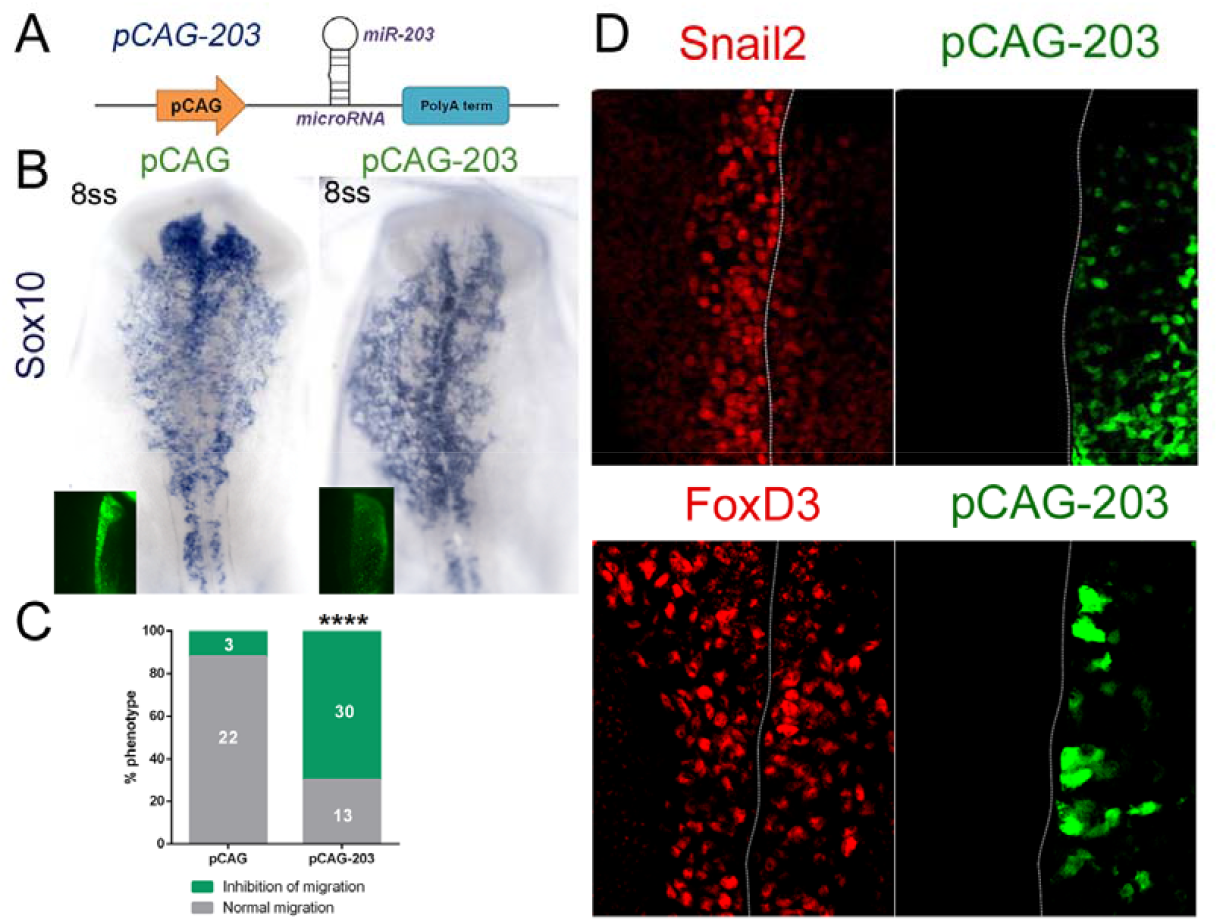
miR-203 overexpression prevents NCC delamination without affecting specification. (A) Scheme of pCAG-203 vector to overexpress miR-203. (B) *In situ* hybridization for *Sox10* showed inhibition of NCC migration in the pCAG-203 injected side, in comparison with the uninjected side of the same embryos and with embryos injected with an empty pCAG vector. (C) Quantitation of the embryos according to their phenotypes (normal migration versus inhibition of migration) was analyzed. Numbers in the graphs represent the numbers of analyzed embryos. ****P < 0,0001 by contingency table followed by X^2^ test. (D) Immunohistochemistry analyzes on miR-203 overexpressing embryos evidenced a reduced expression of SNAIL2 but does not affect the expression of the early NCC specifier FOXD3.

### Loss of miR-203 function causes premature NCC delamination

Given that miR-203 is epigenetically repressed in pre-migratory NCC and overexpression of miR-203 causes defect in their delamination, we next asked if early loss of miR-203 function would result in premature neural crest delamination. To test this possibility we adapted a protocol (Kluiver et al., 2012) to generate a “sponge” vector containing repeated miR-203 antisense sequences (pSmiR-203) to sequester endogenous miR-203 (Fig. 4A). A sponge vector containing a scrambled sequence (pSmiR-scramble) was designed as a control. As predicted, electroporation of pSmiR-203 resulted in premature NCC migration, when compared with the contralateral uninjected side, analyzed by immunohistochemistry for SNAIL2 and FOXD3, or by ISH for *Sox10* (Fig. 4B). Importantly, no difference in timing of delamination was observed after electroporation of the pSmiR-scramble. By categorizing embryos according to their phenotype, premature versus normal migration, we observed a significant difference in percentage of embryos exhibiting premature NCC migration in pSmiR-203 electroporated embryos (Fig. 4C). Notably, reduction of miR-203 function altered not only initiation of NCC delamination, but also shortened the overall length of time during which delamination occurred (Fig. 4D-D’). This is based on the observation that on the side injected with pSmiR-203, all the Sox10+ NCCs have completed their delamination, compared with the contralateral side where many premigratory neural crest cells still persist on the dorsal neural tube. These results clearly confirm that miR-203 controls the temporal regulation of NCC delamination.

**Figure 4:**
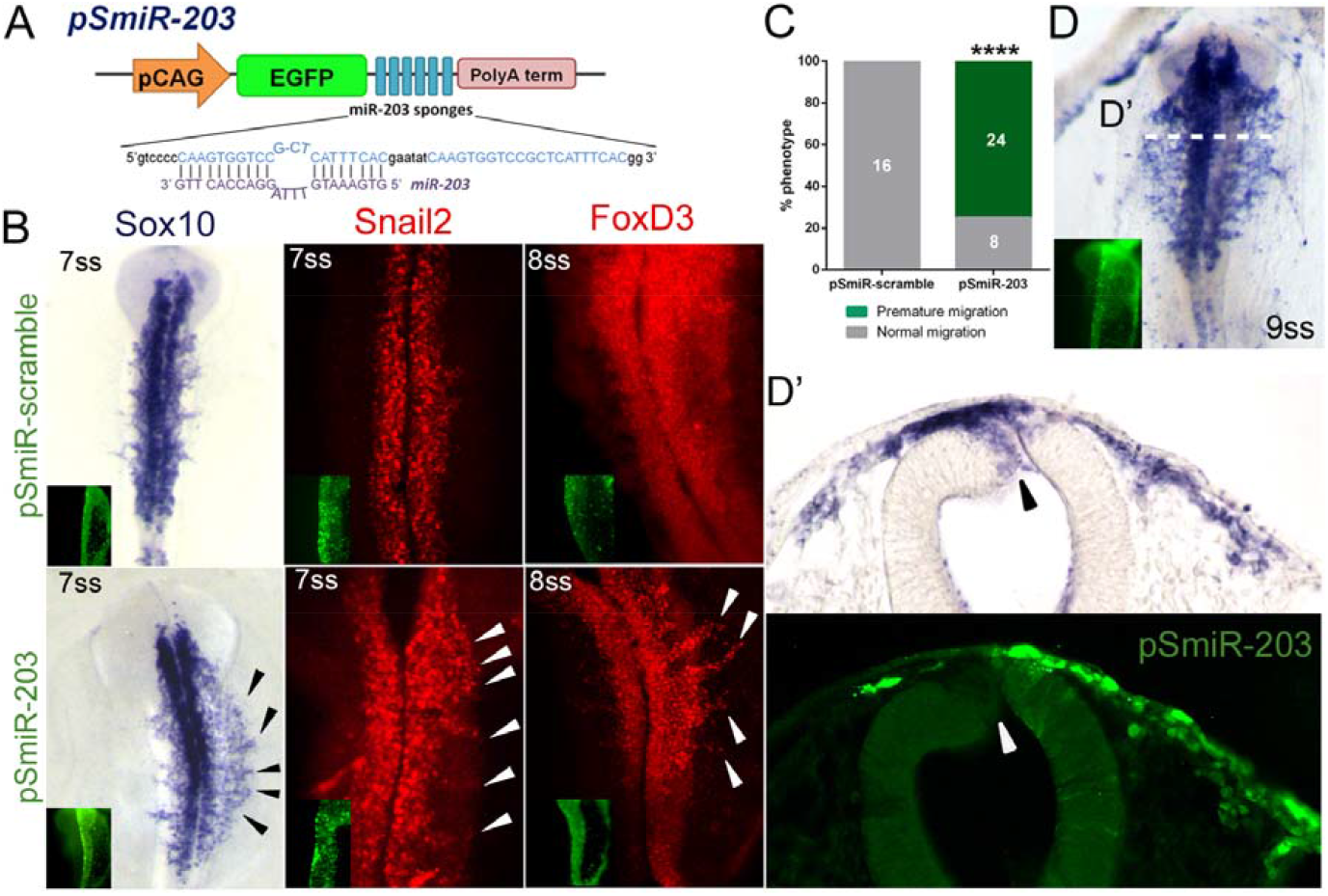
miR-203 sponge vector causes premature NCC delamination. (A) Scheme of pSmiR-203 sponge vector having a bulged miR-203 antisense sequences downstream of the EGFP reporter. (B) Electroporated embryos with pSmiR-203 causes premature migration of NCC evidenced by *in situ* hybridization for Sox10 and immunohistochemistry for SNAIL2 and FOXD3, and compared with the uninjected side of the same embryos or injected with pSmiR-scramble vector. Arrowhead indicated premature migratory neural crest cells. (C) Quantitation of pSmiR-203 or pSmiR-scramble treated embryos according to the observation of premature NCC migration. Numbers in the graph represent the analyzed embryos. ****P<0,0001 by contingency table followed by X^2^ test. (D-D’) Neural crest cells from the sponge injected site complete their delamination in advance compare with the uninjected site where many *Sox10* expressing cells are still on the neural tube (see black arrowhead).

### miR-203 targets the 3’UTRs of Snail2 and Phf12

To test whether *Snail2* and *Phf12* genes are direct targets of miR-203, we designed two-colored sensor vectors in which we cloned, downstream of the pCAG and d4EGFPN, the 3’UTRs containing the wild (pUTR-Snail2/Phf12) or mutated (pUTR-mutSnail2/Phf12) miR-203 binding sites (Fig. 5A). Coelectroporation of these vectors (Fig. 5B) demonstrated that overexpression of miR-203 specifically inhibited d4EGFP_N_ expression when it was fused to the 3’UTRs of *Snail2* and *Phf12* but was uniformly distributed when miR-203 binding sites were mutated (Fig. 5C-D). These results confirm that *Snail2* and *Phf12* are endogenous targets of the same microRNA, miR-203.

**Figure 5:**
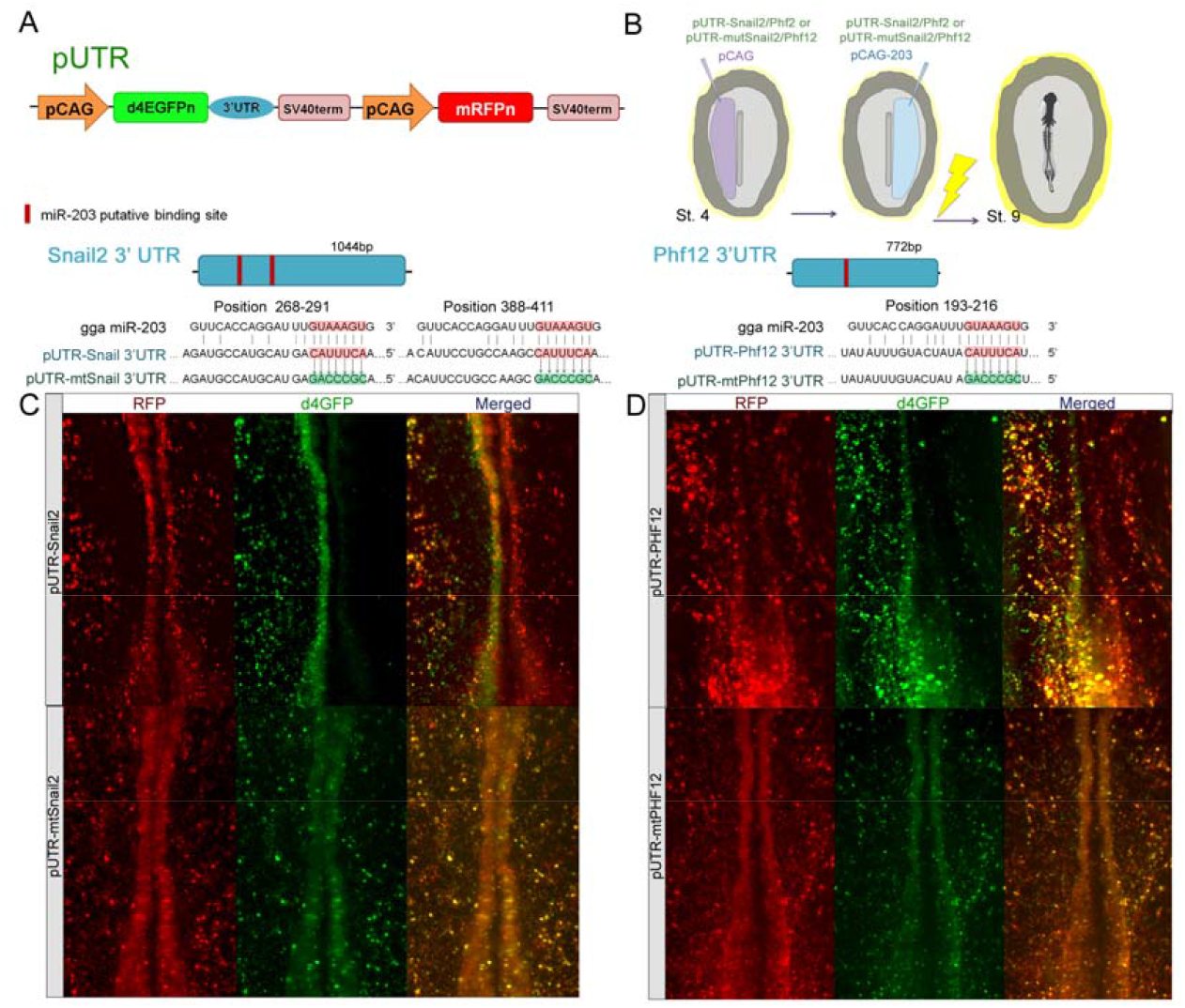
*Snail2* and *Phf12* 3’UTRs are direct targets of miR-203. (A) Scheme of dual colored sensor vector containing wild or mutated (mt) 3’UTRs from Snail2 (pUTR-Snail2) and Phf12 (pUTR-Phf12). pCAG, Chick β-actin promoter; d4EGFP_N_ nuclear-localized destabilized EGFP with a half-life of 4 h; mRFP_N_, nuclear-localized monomeric red fluorescent protein. (B) Diagram of electroporation assays for 3’UTR-sensor experiments. Electroporation of sensor vector containing the 3’UTRs for (C) *Snail2* (pUTR-Snail2) or (D) *Phf12* (pUTR-Phf12) together with miR-203 overexpressing vector (pCAG-203) causes a consistent reduction on the d4EGFPN expression (right side) compared with the control side (left side). Mutation of miR-203-binding sites in the 3’UTRs of *Snail2* (pUTR-mtSnail2) or *Phf12* (pUTR-mtPhf12) caused that most of the electroporated cells were yellow, expressing both d4EGFPN and mRFPN, even when miR-203 is overexpressed (right side).

## Discussion

There is accumulating evidence for the importance of microRNAs in normal development as well as in several diseases, including tumor metastasis. Our study highlights the key role of a single microRNA, miR-203, in regulating the timing to initiation of the EMT program in neural crest cells. The results show that repression of miR-203 occurs via high levels of DNA methylation of the miR-203 locus by the DNMT3B, whose specific activity is directed by SNAIL2. In this scenario, repression of miR-203 is directed by SNAIL2 in a feedback-loop that enables expression of two direct targets of miR-203, *Phf12* and *Snail2,* which in turn are necessary for neural crest delamination. Finally, miR-203 gain-and loss-of-function cause reduction or premature NCC delamination, respectively. Taken together, the results reveal for the first time an epigenetic-miRNA-gene regulatory circuit that controls the timing of neural crest delamination (Fig. 6A). These findings support the idea that a single microRNA may “throw the switch” from an epithelial to a mesenchymal state in the neural crest and thus stabilize the core gene regulatory networks in these two states. There is increasing evidence to suggest that miRNAs often act as fine-tuning regulators rather than as primary gene regulators (Hornstein and Shomron, 2006). Accordingly, we postulate that miR-203 may act by shifting the levels of SNAIL2 and PHF12 to prevent premature NCC delamination. A similar epigenetic-miRNA control of the core transcription factors necessary for EMT (EMT-TFs) has been also described in cancer cells (Wright et al., 2010, Guttilla et al., 2012, Xia and Hui, 2012, Kiesslich et al., 2013, Ding, 2014). Interestingly, some miRNAs and EMT-TFs form a tightly interconnected feedback loop that controls epithelial cell plasticity (Moes et al., 2012, Ding, 2014, Wellner et al., 2009), similar to our observation that SNAIL2 directs the epigenetic repression of miR-203. These feedback-loops provide self-reinforcing signals and robustness to maintain the epithelial or mesenchymal cell state in response to different environmental cues. miR-203 is highly conserved from lamprey to human and appears to be an evolutionary novelty in vertebrates (Heimberg et al., 2010). Our data demonstrate that miR-203 is a key regulator of NCC delamination by a reversible epigenetic-miRNAs. Given that the EMT program is a highly conserved process, involving similar transcription factors in both embryonic and cancer cells, these finding open new avenues for understanding normal and pathological development, as well as tumor metastasis.

**Figure 6:**
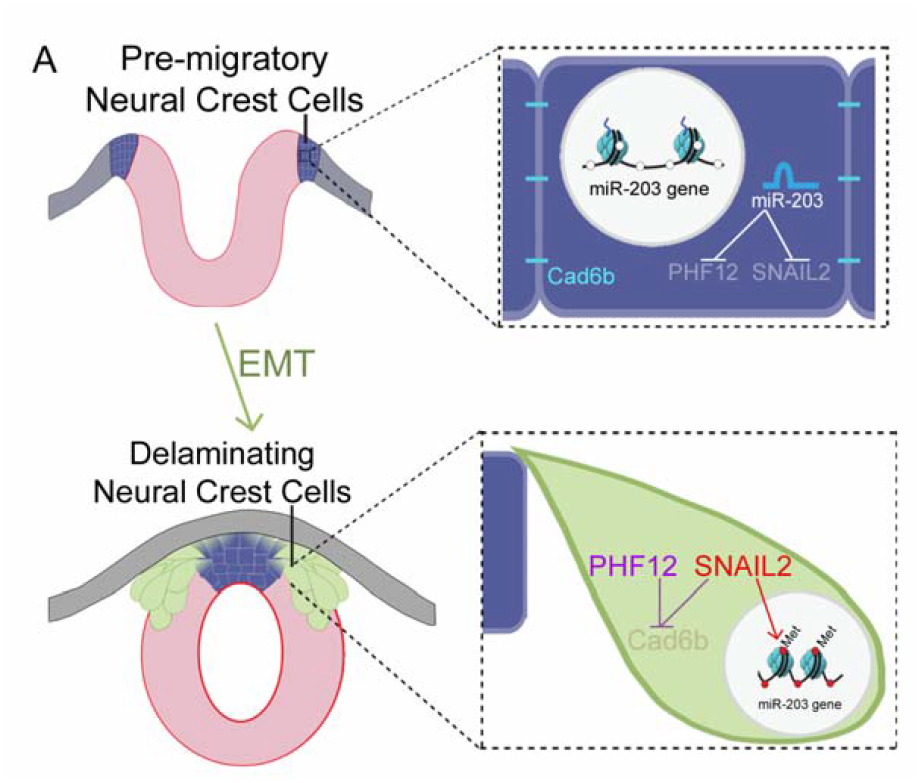
Hypothetical model. Our results show that during NCC specification, miR-203 is highly expressed and preventing the initial accumulation of SNAIL2 and PHF12. Previous to NCC delamination, the accumulation of SNAIL2 causes the epigenetic silencing, mediated by DNMT3B, of miR-203. Then, the lack of miR-203 allows a rapid upregulation of both *Snail2* and *phd12* at the same time, which are necessaries for Cad6b repression at the beginning of the NCC epithelial-to-mesenchymal transition.

## Materials and methods

### RNA preparation and RT-qPCR

RNA was prepared from individual embryos (n=6) using the isolation kit RNAqueous-Micro (Ambion) following the manufacturer’s instruction. The RNA was treated with amplification grade DNaseI (Invitrogen) and then reversed transcribed to cDNA with a reverse transcription kit (SuperScript II; Invitrogen) using stem-loop-miRNA-specifics (Chen et al., 2005) primers and random hexamers. QPCRs were performed using a 96-well plate qPCR machine (Stratagen) with SYBR green with ROX (Roche). Normalization controls genes for qPCR were: for miR-203, miR-16 (Lardizabal et al., 2012) and for *Snail2* and *Phf12, Hprt1* (Simoes-Costa and Bronner, 2016). For a complete list of primer see table S3.

### Bisulfite sequencing

Samples were obtained by dissecting of 9 embryos at stage 6ss, for premigratory NCC (PM-NCC), dorsal neural tube, and ventral neural tube (NT). In addition, we dissected 13 embryos at stage 11-13ss to obtain the migratory NCC (M-NCC). For the morpholino-mediated loss of DNMT3A, DNMT3B and SNAIL2 experiment, eight dorsal neural tubes from the injected and uninjected sides were dissected. For a complete list of morpholinos see table S3. All the tissues were lysed and bisulfite-converted with the EpiTect Plus Bisulfite Conversion Kit (Qiagen) following the manufacturer’s instructions. The regulatory regions of miR-203 were amplified by using two sets of nested primers (see table S3) from the bisulfite-converted DNA. The obtained products were gel-purified and cloned into the pGEM-T Easy Vector (Promega). Individual clones were sequenced and analyzed.

### Electroporation

Chicken embryos were electroporated at stage 4-5 using previously described techniques (Sauka-Spengler and Barembaum, 2008). The vectors and morpholinos were injected by air pressure using a glass micropipette and platinum electrodes were placed vertically across the chick embryos and electroporated with five pulses of 5.5 V for 50 ms at 100 ms intervals. Embryos were cultured in 0.5 ml albumen in tissue-culture dishes until the desired stages. Embryos were then removed and fixed in 4% PFA and used for immunohistochemistry or *in situ* hybridization.

### In situ hybridization

Whole-mount chick *in situ* hybridization for mRNAs and for microRNA was performed as described previously (Acloque et al., 2008, Darnell et al., 2006). LNA probe for miR-203 used in the assay were obtained from Exiqon and DIG-labelled by using the DIG oligonucleotide 3’ end labeling kit (Roche). After ISH, some embryos were fixed in 4% PFA in PBS, washed, embedded in gelatin, and cryostat sectioned at a thickness of 14–16 μm. They were photographed using the NIS-Elements Advanced Research software (Nikon) with an Eclipse E600 microscope (Nikon) and processed using Photoshop CS3 (Adobe).

### Immunohistochemistry

Whole-mount chick immunohistochemistry was performed as described previously (Taneyhill et al., 2007). Briefly, embryos were fixed for 15 min in 4% PFA and then permeabilized and blocked in TBS (500 mM Tris-HCl, pH 7.4, 1.5 M NaCl, and 10 mM CaCl_2_) containing 0.1% Triton X-100 (TBS-T) and 5% FBS for 60 min at room temperature. Primary antibodies were diluted in TBS-T/FBS and incubated overnight at 4°C. Primary antibodies used were mouse anti-Snail2 (1:100), (1:100; supplied by the Developmental Studies Hybridoma Bank), and rabbit anti-FoxD3 (1:300; gift of P. Labosky, Vanderbilt University Medical Center, Nashville, TN). Secondary antibodies used were goat antimouse and anti-rabbit Alexa Fluor 594 (1:500; all obtained from Molecular Probes) diluted in TBS-T/FBS and incubated for 45 min at room temperature. All washes were performed in TBS-T at room temperature. Some embryos were subsequently embedded in gelatin, cryostat sectioned at 12-16μm, photographed using the NIS-Elements Advanced Research software (Nikon) with an Eclipse E600 microscope (Nikon) and processed using Photoshop CS3 (Adobe).

### MicroRNA sponge generation

Oligos designed to generate miR-203 sponge were ordered phosphorylated and PAGE purified at a 100 nmol scale (see table S2) and dissolved to 50 mM in STE buffer (100 mM NaCl, 10 mM Tris/HCL, 1 mM EDTA, pH 8.0). Sense and antisense oligos were mixed at a 1:1 ratio and annealed by incubation at 100°C for 10 minutes followed by slow cooling. The “sponge” vector was generated following previously described protocols (Kluiver et al., 2012).

## Acknowledgments

We thank Dr. Xinwei Cao for the two-colored sensor vector, and Dr. Andrew Pollok for the advices on the design of sponge vector.

## Competing interests

The authors declare no conflicts of interest

## Funding

This work was supported by the Fogarty International Center of the NIH (R21TW011224 to MEB and PHS-M) and the Agencia Nacional de Promoción Científica y Tecnológica (PICT 2016-0747 to PHS-M).

## Author contributions

ESV, MEB and PHS-M conceived, designed and analyzed the experiments. ESV performed all the experiments. ESV, MEB and PHS-M wrote the manuscript.

## Supplemental materials

**Figure S1.**
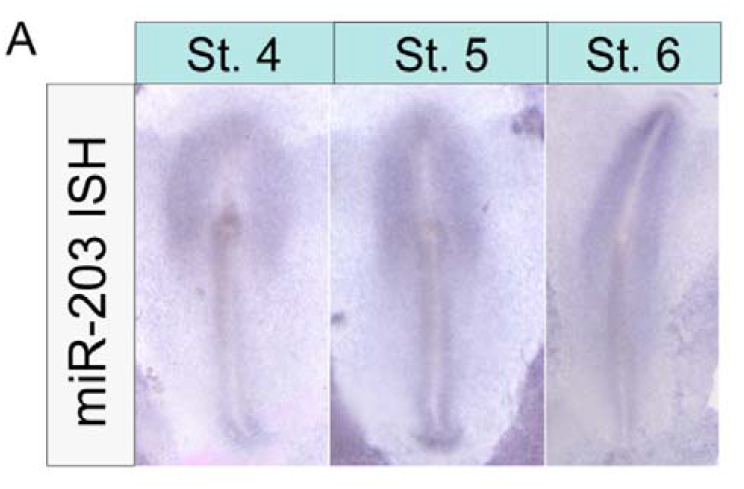
(A) Whole-mount *in situ* hybridization using DIG-labeled LNA probes (Exiqon) against miR-203 at early chick developmental stages.

**Figure S2:**
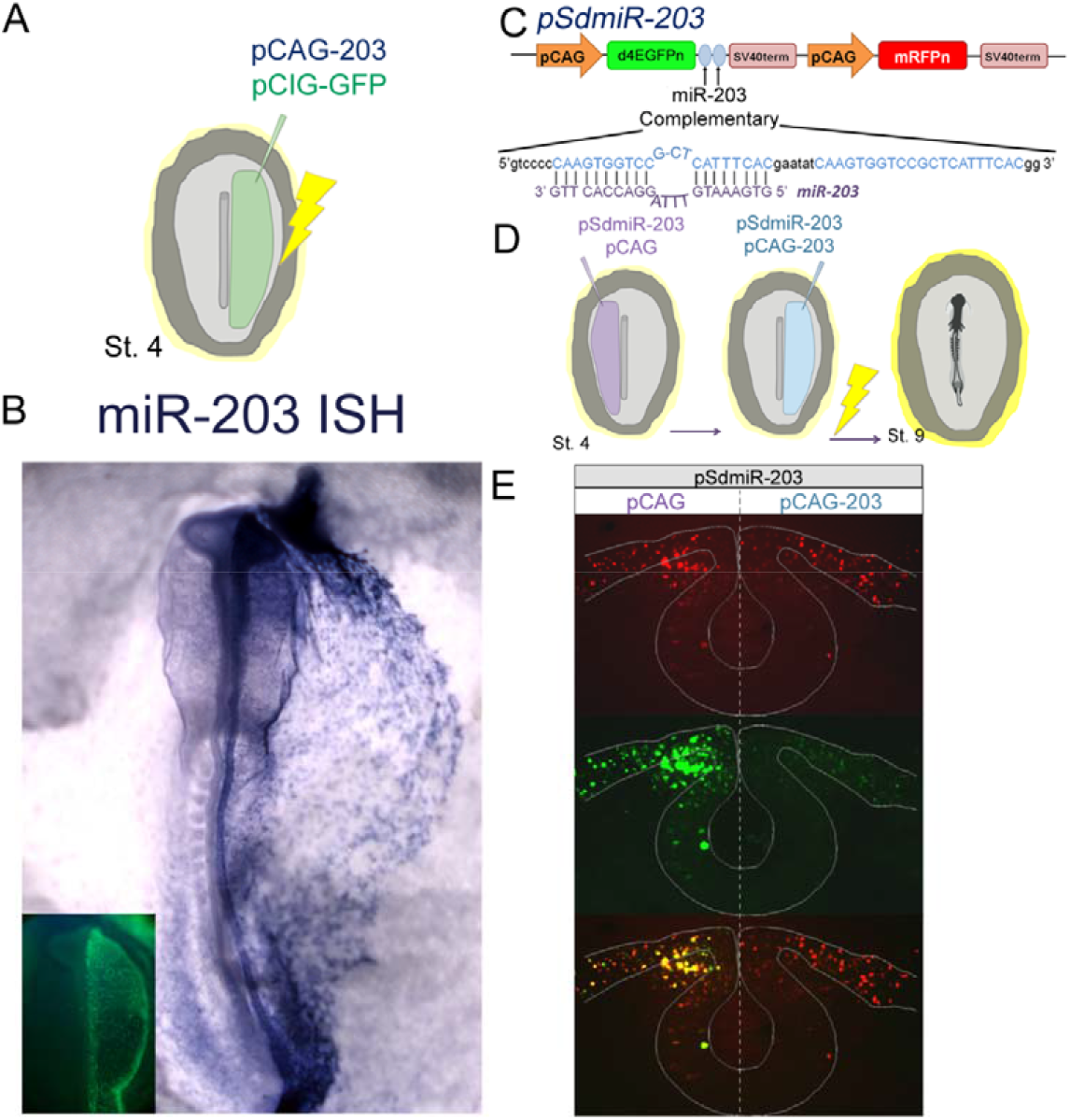
(A) Diagram of electroporation assay for gain-of-function experiments. pCAG-miR-203 does not have a fluorescent marker, for this reason, was co-electroporated with a fluorescent vector that express GFP downstream the CAG promoter. We injected the vectors in the right side of the embryos at stage 4. Following injection, embryos were electroporated and cultured until stage 9. (B) Electroporation of pCAG-203 vector (right side) successfully overexpress a mature miR-203 evidenced by *in situ* hybridization using LNA probes. (C) Schematic drawing of the miRNA dual-sensor vector (pSdmiR-203) containing two copies of complementary sequences to the mature miR-203. (D) Illustration of bilateral electroporation assay to evaluate if pCAG-203 express a functional miR-203. (E) Co-electroporation pSdmiR-203 and the empty pCAG vector (left side) caused that most of the cells are yellow because of the expression of both green and red reporters. Whereas, coelectroporation pSdmiR-203 and pCAG-203 vector (right side) caused that most of the cells are only red, because of the strong repression of the green reporter. pCAG, Chick β-actin promoter; d4EGFPN nuclear-localized destabilized EGFP with a half-life of 4 h; mRFP_N_, nuclear-localized monomeric red fluorescent protein.

**Supplementary table 1:**
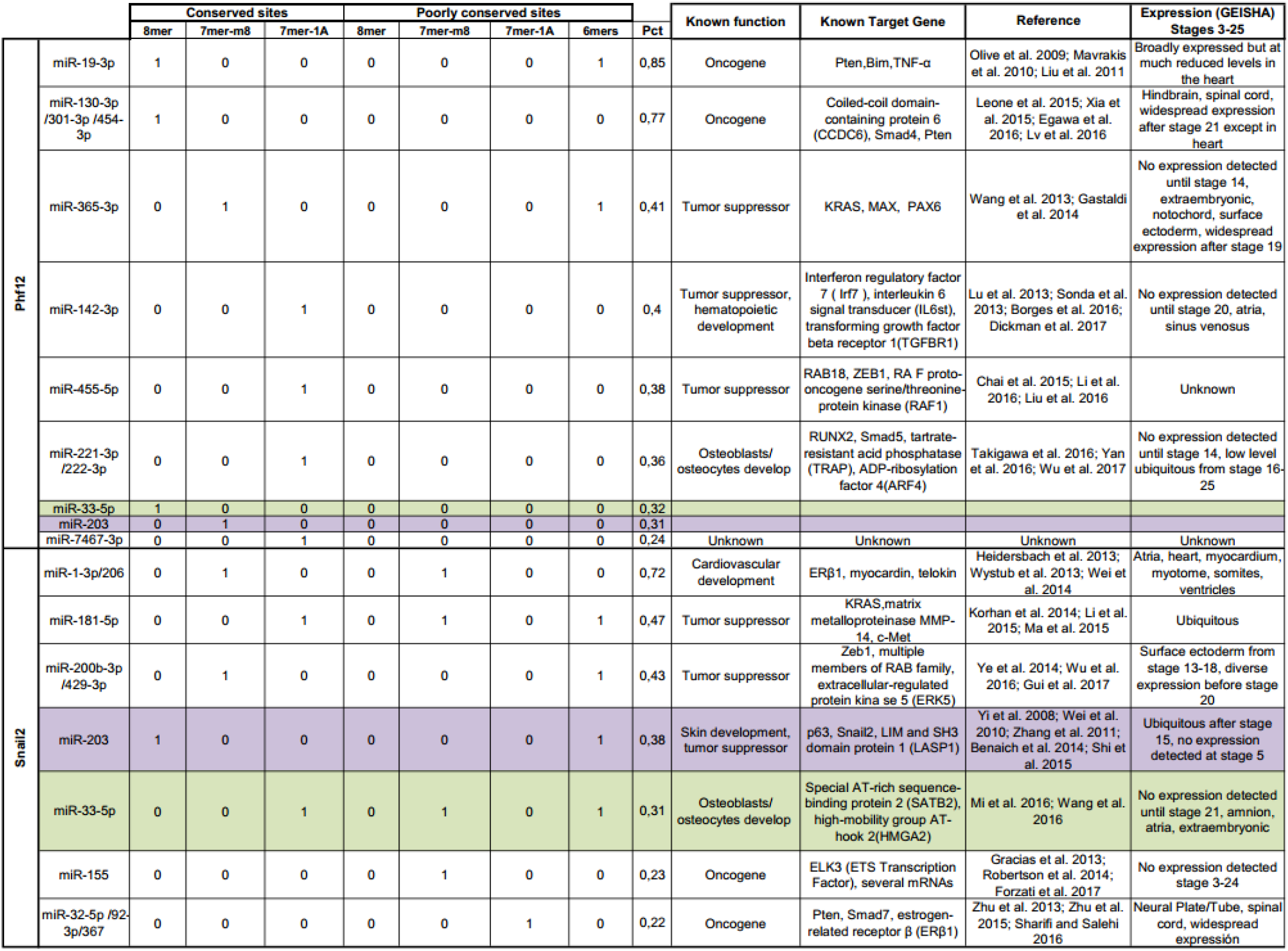
*In silico* analyzes of conserved and poorly conserved microRNA-binding sites on Snail2 and Phf12 3’UTRs (TargetScan), their known functions, demonstrated targets, and chick expression (GEISHA).

**GEISHA microRNAs expression in chick development** http://geisha.arizona.edu/geisha/quick_search.jsp?table=mir_miR-1 (Wei et al., 2014, Wystub et al., 2013, Heidersbach et al., 2013)

**miR-19** (Olive et al., 2009, Mavrakis et al., 2010, Liu et al., 2011)

**miR32-5p/92-3p/367** (Zhu et al., 2013a, Zhu et al., 2015, Sharifi and Salehi, 2016)

**miR-33-5p** (Mi et al., 2016, Wang et al., 2016)

**miR130-3p/301-3p/454-3p** (Leone et al., 2015, Xia et al., 2015, Lv et al., 2016, Egawa et al., 2016)

**miR-142** (Lu et al., 2013, Borges et al., 2016, Sonda et al., 2013, Dickman et al., 2017)

**miR-155** (Robertson et al., 2014, Gracias et al., 2013, Forzati et al., 2017)

**miR-181-5p** (Ma et al., 2015, Li et al., 2015, Korhan et al., 2014)

**miR-200b-3p/429-3p** (Gui et al., 2017, Ye et al., 2014, Wu et al., 2016)

**miR-203**(Wei et al., 2010, Yi et al., 2008, Zhang et al., 2011, Shi et al., 2015, Benaich et al., 2014)

**miR-221-3p/222-3p** (Yan et al., 2016, Takigawa et al., 2016, Wu et al., 2017)

**miR-365-3p** (Wang et al., 2013, Gastaldi et al., 2014)

**miR-455-5p** (Liu et al., 2016, Li et al., 2016, Chai et al., 2015)

---

**Supplementary table 2:** Results obtained with the Jaspar 2018 (http://jaspar.genereg.net/) for SNAIL2 binding site in the tentative promoter of miR-203. High binding sites (>9) are mapped in figure 2A. We also show the sequence analyzed in Jaspar 2018.

| Matrix ID | Name | Score | Relative score | Sequence ID | Start | End | Strand | Predicted sequence |
| --- | --- | --- | --- | --- | --- | --- | --- | --- |
| MA0745.1 | SNAI2 | 12,6931 | 0.999707221414 | miR_203_tentative_promoter | -1162 | -1153 | + | AACAGGTGC |
| MA0745.1 | SNAI2 | 9,71729 | 0.94034684341 | miR_203_tentative_promoter | -1552 | -1543 | + | GGCAGGTAC |
| MA0745.1 | SNAI2 | 8,48966 | 0.915858769456 | miR_203_tentative_promoter | -1089 | -1080 | - | CACAGGTTG |
| MA0745.1 | SNAI2 | 7,98092 | 0.905710714071 | miR_203_tentative_promoter | -1273 | -1264 | - | ATCAGGTTG |
| MA0745.1 | SNAI2 | 4,63965 | 0.839061031445 | miR_203_tentative_promoter | -1599 | -1590 | - | TGCATGTTT |
| MA0745.1 | SNAI2 | 3,94346 | 0.825173896668 | miR_203_tentative_promoter | -1527 | -1518 | + | TCAAGGTGT |
| MA0745.1 | SNAI2 | 3,5435 | 0.817195747557 | miR_203_tentative_promoter | -1199 | -1190 | - | TGGAGGTTG |
| MA0745.1 | SNAI2 | 3,51455 | 0.816618237138 | miR_203_tentative_promoter | -1293 | -1284 | + | AGAAAGTGA |
| MA0745.1 | SNAI2 | 3,24005 | 0.811142584031 | miR_203_tentative_promoter | -1082 | -1071 | + | TGCCAGTGC |

>miR\_203\_tentative\_promoter Chr 5:50767590-50768212

**Supplementary table 3:**
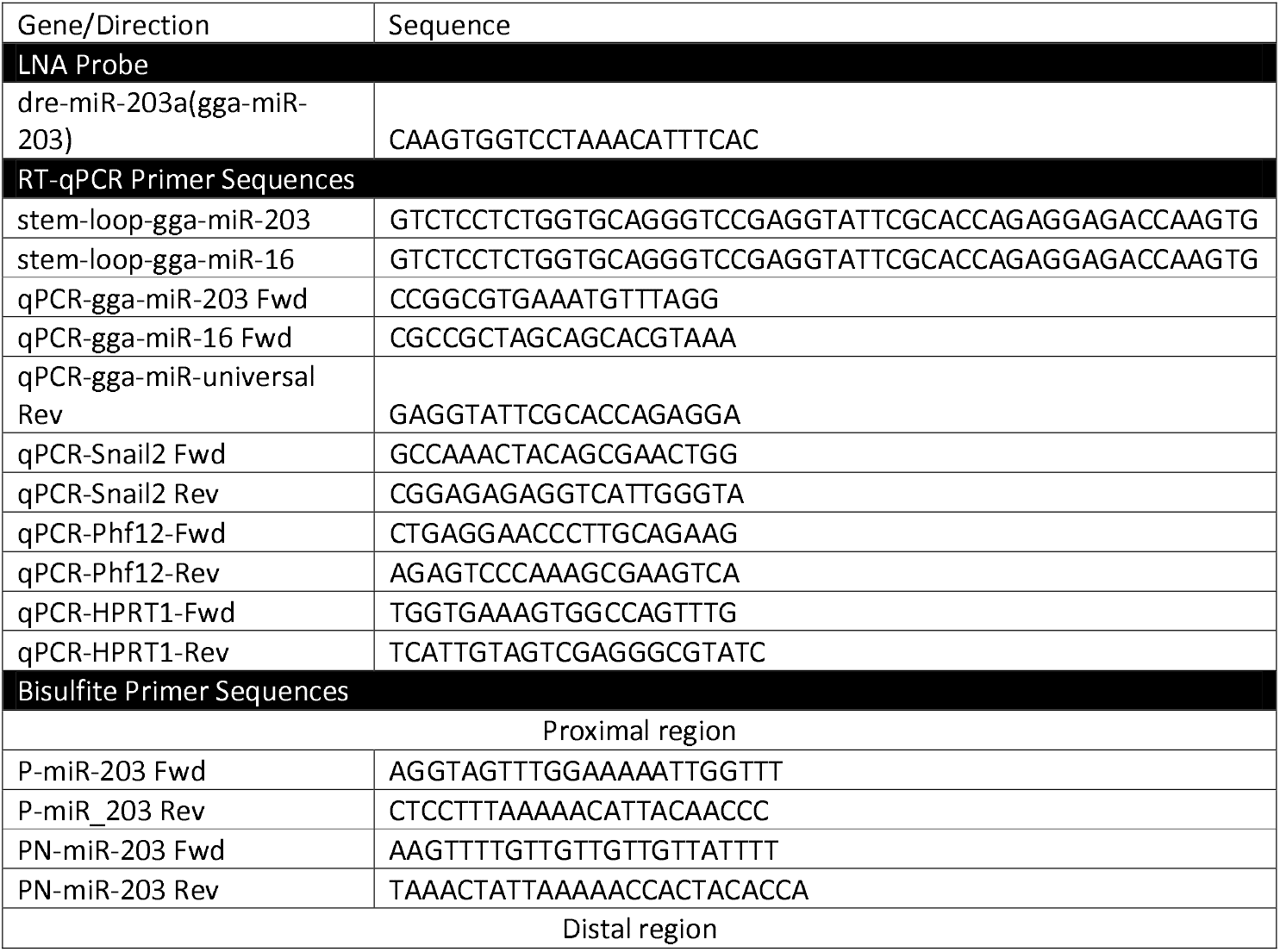
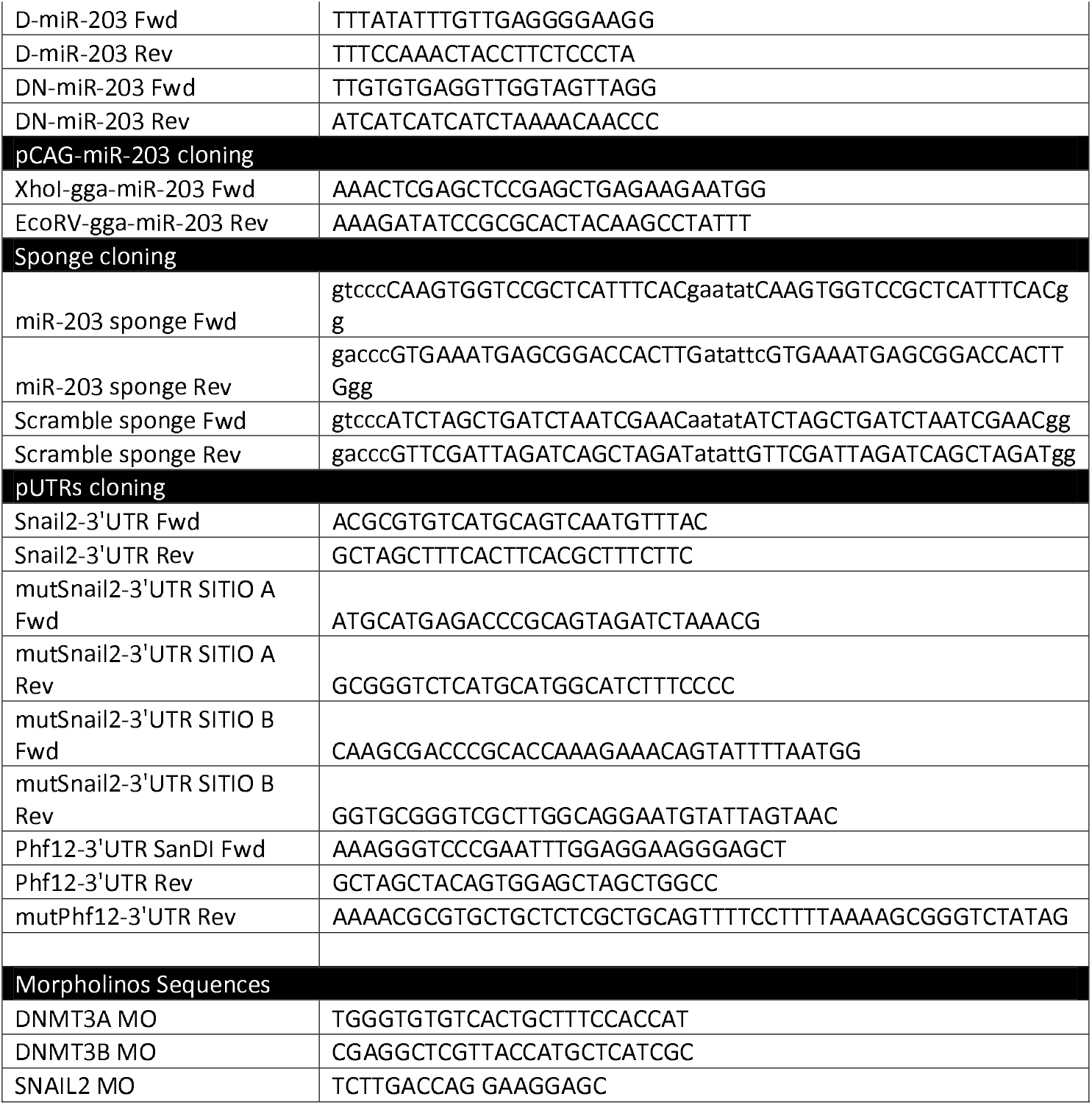
Complete list of utilized primers, LNA probe, and morpholinos

